# Convergent recruitment of *Amh* as the sex determination gene in two lineages of stickleback fish

**DOI:** 10.1101/2022.01.26.477894

**Authors:** Daniel L. Jeffries, Jon Mee, Catherine L. Peichel

## Abstract

Sex chromosomes vary greatly in their age and levels of differentiation across the tree of life. This variation is largely due to the rates of sex chromosome turnover in different lineages; however, we still lack an explanation for why sex chromosomes are so conserved in some lineages (e.g. Mammals, Birds) but so labile in others (e.g. Fish, Amphibians). Here we add to the information on sex chromosomes in stickleback, a valuable model lineage for the study of sex chromosome evolution, by identifying the sex chromosome and a strong candidate for the master sex determination gene in the brook stickleback, *Culaea inconstans*. Using whole genome sequencing of wild-caught samples and a lab cross, we identify *AmhY*, a male specific duplication of the gene *Amh*, as the candidate master sex determination gene. *AmhY* resides on Chromosome 20 in *C. inconstans* and is likely a recent duplication, as both *AmhY* and the sex linked region of Chromosome 20 show little sequence divergence. Importantly, this duplicate *AmhY* represents the second independent duplication and recruitment of *Amh* as the sex determination gene in stickleback and the eighth example now known across teleosts. We discuss this convergence in the context of sex chromosome turnovers and the role that the *Amh*/*AmhrII* pathway, which is crucial for sex determination, may play in the evolution of sex chromosomes in teleosts.

## Introduction

Upon acquiring a master sex determination gene (MSD), sex chromosomes are set on an evolutionary trajectory different to that of the rest of the genome. In many cases recombination is reduced or suppressed entirely in the vicinity of the MSD, which opens the door to the accumulation of deleterious mutations due to Hill-Robertson interference and Muller’s ratchet. Given enough time this process can lead to loss of gene function on the sex chromosomes and eventually to hetermorphic sex chromosomes (Charlesworth *et al*., 2005), typified by those of Mammals and Birds (Bachtrog *et al*., 2014).

In some taxa, however, sex chromosomes escape this process via sex chromosome turnovers, the swapping of the chromosome used for sex determination. When this occurs, sex chromosome differentiation is reset (Vicoso, 2019) and, as such, lineages with labile sex determining systems often have homomorphic, undifferentiated sex chromosomes (Jeffries *et al*., 2018). Sex chromosome turnovers can, therefore, be seen as one of the most impactful processes in the evolution of sex chromosomes and indeed the genome. However, we still lack a good understanding of what drives turnovers. One theory is that the accumulation of deleterious mutations in a sex linked, non-recombining region should favour a transition to a new sex determination gene as a means of purging the genome of mutation load (Blaser *et al*., 2014). Alternatively, a transition to a new sex chromosome may be favoured if it harbours a sexually antagonistic gene which, when linked to a new MSD, provides a fitness increase to both sexes (van Doorn & Kirkpatrick, 2007, 2010). Finally, turnovers may occur simply via drift (Saunders *et al*., 2018). Understanding the relative importance of these drivers, and other factors that may contribute to sex chromosome evolution and transitions is essential to explain the diversity and distribution of sex determining systems in nature. However, we still lack sufficient empirical evidence from lineages with young and labile sex determining systems with which to test the above theories.

Sticklebacks (Teleostei: Gasterosteidae) are one such lineage, possessing a diverse set of sex chromosome systems across their phylogeny (Ross *et al*., 2009; Dixon *et al*., 2018; Natri *et al*., 2019; Peichel *et al*., 2020; Sardell *et al*., 2021). Species of the *Gasterosteus* genus share a relatively well conserved XY sex determining system located on Chromsome 19, which harbours a strong candidate for sex determination in *Gasterosteus, AmhY*, a Y-specific duplicate of the ancestrally autosomal gene *Amh*. This sex chromosome is estimated to have evolved approximately 22 million years ago (Peichel *et al*., 2020) and is heteromorphic (Ross & Peichel, 2008), with considerable loss of genes on the Y (Peichel *et al*., 2020; Sardell *et al*., 2021). However, in two *Gasterosteus* species, *G. nipponicus* and *G. wheatlandi*, independent Y-autosome fusion events have created neo sex chromosomes which became sex linked within the past two million years (Kitano *et al*., 2009; Ross *et al*., 2009). In the genus *Pungitius*, chromosome 19 is not known to be involved in sex determination, instead, Chr 12 determines sex in *P. pungitius* (Ross *et al*., 2009; Shapiro *et al*., 2009; Rastas *et al*., 2015; Natri *et al*., 2019). However, despite interrogation of high quality genomic datasets for both *P. sinensis* and *P. tymensis*, no sex chromosome has yet been identified for these species (Dixon *et al*., 2018). Similarly, no sex chromosome could be identified in either *Apeltes quadracus* or *Culaea inconstans* using either cytogenetic techniques or genetic mapping with a small number of markers (Ross *et al*., 2009). This suggests that the sex chromosomes of these species have little divergence between gametologs, making it likely that recent turnovers have occurred in at least some of these species. The sex chromosomes of stickleback therefore include old strata that have undergone degeneration, newly-formed and weakly differentiated strata on neo-sex chromosomes, and likely several new and undifferentiated sex chromosomes created by recent turnovers. As such, stickleback represent an invaluable system to test several facets of sex chromosome evolution theory.

Here, we add to the information of sex chromosome systems in sticklebacks by identifying the sex chromosome and a strong candidate for the sex determination gene in the brook stickleback, *Culea inconstans*. We show that *C. inconstans* has undergone a sex chromosome turnover to a chromosome not previously known to be used in stickleback. This turnover seems to have been driven by a duplication and translocation of the well known sex determination gene, *Amh*, which is independent of the duplication of this gene in the *Gasterosteus* lineage. Thus, although the turnover involves a novel chromosome pair, the sex determination gene itself represents an example of convergence both in terms of function and the mode of turnover, and may provide clues as to why sex chromosomes in some lineages are so dynamic.

## Methods

### Sample collection and sequencing

In this study we examined two sample sets of the brook stickleback, *C. inconstans*. The first and largest is comprised of wild-caught individuals from a single population in Shunda Lake, Alberta, Canada (UTF-8 encoded WGS84 coordinates: 52.453899 latitude, -116.146192 longitude). We collected a total of 84 samples in June of 2017 and 2019 using unbaited minnow traps (5 mm mesh). Samples were collected under a fisheries research license issued by the Government of Alberta. Collection methods and the use of animals in research was approved by the Animal Care Committee at Mount Royal University (Animal Care Protocol ID 101029 and 101795). We identified 46 males and 38 females at the site of capture by examining gonads, observing the extrusion of eggs, and by noting the presence of nuptial colouration in males. DNA was extracted using Qiagen DNEasy Blood and Tissue kits. DNA samples were sent to Genome Québec for shotgun DNA library preparation using an NEB Ultra II kit. Paired-end sequencing (150bp) was performed alongside other libraries; the samples in this study therefore received approximately one lane of Illumina HiSeqX (40 samples collected in 2017) and half of a NovaSeq6000 lane with an S4 flow cell (remaining 44 samples collected in 2019).

The second sample set consists of a single F1 lab cross between a female from Fox Holes Lake, Northwest Territories, Canada and a male from Pine Lake, Alberta, Canada; this cross was previously genotyped with a limited set of microsatellite markers (Ross *et al*., 2009). DNA was isolated from fin tissue using phenol-chloroform extraction followed by ethanol precipitation. Four Nextera sequencing libraries were prepared: one using DNA from the mother, one using DNA from the father, one using DNA pooled from 16 daughters, and one using DNA pooled from 14 sons. Paired-end sequencing was performed on an Illumina NovaSeq SP flow cell for 300 cycles at the University of Bern Next Generation Sequencing Platform.

### Data pre-processing and SNP calling in wild-caught C. inconstans

Sequencing of the 84 Shunda lake stickleback yielded an average of 24.8 million read pairs per sample. Sequence quality was checked using FastQC and an average of 0.59% (± 0.13%) read pairs per sample were dropped during adapter and quality trimming using Trimmomatic v.0.36 (Bolger *et al*., 2014). Trimmed reads were then aligned using BWA-mem v.0.7.17 (Li & Durbin, 2009) with default alignment parameters to a genome assembly of a *P. pungitius* male (Varadharajan *et al*., 2019) as it is the most closely related reference genome to *C. inconstans* (21.16 - 24.30 MYA) (Rabosky *et al*., 2013, 2018; Betancur-R *et al*., 2015; Sanciangco *et al*., 2016; Guo *et al*., 2019). Alignment files were then processed with samtools v.1.10 and an average of 16.4% (± 7.0%) read pairs were then marked as PCR duplicates and removed using picard-tools v.2.21.8. Remaining aligned reads resulted in an average read depth of 7.20 (± 2.14) for each sample; however, coverage was highly variable along the genome, with peaks of coverage of over 1,500 reads in places, almost certainly driven by repeats.

Variant calling was performed using the Genome Analysis Toolkit (GATK) v.4.1.3.0 following the GATK best practices pipeline (DePristo *et al*., 2011) and resulted in 36,237,609 variants before filtering. Comparison of these variants to Hardy-Weinberg expectations revealed a large number of variants with an excess of heterozygosity, likely as a result of repetitive regions. The full variant call sets for the wild-caught samples were then filtered using VCFtools v0.1.15 (Danecek *et al*., 2011) to retain single nucleotide polymorphisms (SNPs) with the following attributes: within samples or pools, genotypes were retained if they had a minimum depth of 10 reads (--minDP 10) and a minimum genotype quality score of 30 (--minGQ 30). Across samples, loci were kept if they had a maximum mean depth across all samples of 200 or lower (max-meanDP 200), less than 30% missing samples after genotype filters (--max-missing 0.3), and a minor allele frequency greater than 0.01 (--maf 0.01). These filtering criteria reduced the call set to 249,485 variants, which were used for all analyses of the wild-caught dataset below. However many loci showing excess heterozygosity persisted in the dataset, and were not filtered further, as this would likely remove signals of sex linkage (Fig. S1).

### Identification of sex-linked regions of the genome in wild-caught C. inconstans

Sex-linked genome regions typically exhibit two features which can be used to identify them using genomic data analysis. The first is that they often lose or gain segments of DNA on only one of the sex chromosomes. Such regions produce a read depth difference among the sexes in sequencing data reflecting their copy number in the genome. For instance, an X-specific region will have roughly 2n coverage in XX females, and only 1n coverage in XY males, resulting in a ratio of male:female read depth of around 1:2. Secondly, sex-linked regions accumulate sequence differences between the sex chromosomes. This often results in variants which are specific to the sex-limited chromosome, which leads to an increase in SNP density and heterozygosity in sex-linked regions in the heterogametic sex, relative to the homogametic sex. It is generally observed that small mutational differences accumulate on sex chromosomes in the early stages of their differentiation, and large loss or gain of DNA sequence is typically a sign of an older sex linked region. Here, we used both read depth and heterozygosity based approaches for assessing sex linkage across the genome in the wild-caught dataset from Shunda Lake.

For the read depth analysis, Deeptools v.2.5.4 was used to calculate coverage per sample across the genome in 1kb windows, normalised by the average number of reads per kilobase mapped (RPKM) (script 11, appendix 1). Normalised coverages for each window were then averaged for each sex and the mean male depth per window was then divided by that of females and plots were smoothed using a rolling average over 10 windows (see Jupyter notebook JN_02, appendix 1).

We then assessed genotypes for patterns of heterozygosity consistent for sex linkage. SNP calling resulted in a large number of loci with excess heterozygosity, most likely due to reads from multiple repeat copies in *C. inconstans* aligning to a single (likely collapsed) repeat locus in the *P. pungitius* assembly (See above). However, repeats are common in sex-linked genomic regions and therefore likely to contain signals of sex linkage, some of which can be salvaged (i.e. old repeat copies which are unique enough for robust assembly and alignment). Thus, we opted not to mask repeats in the *P. pungitius* genome prior to read alignment. Instead, we used a novel test for the association of heterozygosity at each locus with sex. For each locus, we calculated the probability of the observed pattern of heterozygotes across males and females occurring by chance using a non-sequential random draw without replacement which takes into consideration the number of samples of each sex called at a given locus:

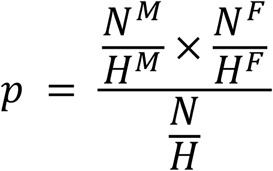

Where N is the total number of samples called, N^M^ is the number of males called, N^F^ is the number of females called, H is the total number of heterozygotes, H^M^ is the number of male heterozygotes and H^F^ is the number of female heterozygotes. The resulting p-values suffer from multiple testing. However, much like in genome wide association studies, the correction is not straightforward as genetic linkage between loci in close proximity to each other violates the multiple testing assumption that each test is independent. Here we avoided this issue by not invoking any threshold for “significant” sex linkage. Instead we simply use our calculated p-values as a relative measure of the extent of sex linkage. For reference, absolute sex linkage of a locus (i.e. heterozygous in all 46 males and homozygous in all 38 females) would yield *p* = 8.6 × 10^−25^ (log2(*p*) = -79.9), while, for a scenario with 20 heterozygotes evenly distributed across the sexes (11 male heterozygotes and 9 female), *p* = 0.2 (log2(*p*) = -2.3) (JN_03, Appendix 1).

### Locating AmhY in the C. inconstans genome

Male coverage patterns suggested that there is an additional sex-linked copy of *Amh* in the *C. inconstans* genome, implying that the duplicate must exist only on the Y chromosome (see Results). If a Y-specific copy of *Amh* exists, then, as an artifact of the read alignment to a genome with only a single copy of *Amh*, any mutations that have arisen in the Y copy since the duplication will be observed as male-specific SNPs within the Chromosome 08 copy of the gene. However, the Y-linked alleles at these sites should show half the coverage (1n) of the autosomal alleles (2n). Thus, to test the hypothesis of a Y-linked duplication of *Amh*, we compared allelic read depth ratios at the sex-linked SNPs located in the *Amy* gene and compared them to the rest of the SNPs throughout the genome in the wild-caught dataset, using a slightly modified implementation of HDplot (McKinney *et al*., 2017) (JN_04, Appendix 1).

We then asked: where in the genome does the sex-linked duplicate of *Amh* (and thus the Y chromosome) reside? Patterns of sex-biased heterozygosity identified two regions of the *P. pungitius* reference genome with signs of sex linkage: *Amh* on Chromosome 08 and Chromosome 20 (see Results). However it is not possible to have two strongly sex linked regions in a genome. Polygenic sex determination systems exist, but complete sex linkage signal would not be observed at either locus, which rules out this possibility in the current study. We therefore hypothesised that the two regions showing sex linkage are an artifact of the alignment to the *P. pungitius* genome assembly, and that, in *C. inconstans*, these sex-linked loci lie in a single region. The most parsimonius scenario is that the duplicated *Amh* copy resides in the region of Chromosome 20 showing sex linkage. However, there were also regions that showed signs of Y-specific duplications in this region of Chromosome 20 (see Results), raising the possibility that this region might also have duplicated, and that the sex determining region in *C. inconstans* may lie on a different chromosome altogether.

To test these competing theories, we first called structural variants in every wild-caught sample using DELLY v0.7.8 (Rausch *et al*., 2012) and screened these variants for any translocations between the regions of sex linkage on Chromosomes 08 and 20, and for any variants showing patterns of sex linkage. Second, we used Abyss v2.0.2 (with default parameters) to produce a *de novo* contig assembly of a pool of raw reads from the three highest coverage *C. inconstans* males, equating to approximately 100x coverage of the genome. The aim of this approach was to produce contigs that included either the autosomal Chromosome 08 copy of *Amh*, or the sex-linked *AmhY* flanked by regions syntenic to the sex linked region of Chromosome 20.

Lastly, we analysed the whole genome resequencing of the lab cross consisting of individually sequenced parents, a female offspring pool, and a male offspring pool. As this cross represents only 30 separate meioses (i.e. 30 offspring) in the father, there should only have been on the order of 30 crossover events between the X and the Y chromosome. This design should thus result in large linkage blocks along the genome, making it much easier to identify regions of the genome which are inherited in a sex-linked fashion.

These sequence data were processed using the same procedures as for the wild-caught data: data were quality checked and trimmed using fastQC and trimmomatic, resulting in 177.6 million retained reads for the father, 168 million for the mother, 160.8 million for the male offspring pool, and 75.7 million for the female offspring pool. These reads were aligned to the *P. pungitius* reference assembly, and duplicates were marked using picard-tools v.2.21.8. We called variants in the parents using bcftools v1.10, which resulted in 19.7 million unfiltered variants in the male and 19.4 million in the female. To call variants in the pooled sequencing data, we used samtools v.1.10 to create an mpileup file which was then converted to allele frequencies using MAPGD v0.4.40 (Lynch *et al*., 2014). Variants were retained (using a custom python script JN_05, Appendix 1) only if they were present in the father, mother, male pool and female pool, and if read depth in the parents was greater than 10 and parental genotype quality was greater than 30. To visualise the data, we plotted female - male allele frequencies along the genome. Sex-linked regions in which females are homozygous and males are heterozygous should show a female - male frequency of close to 0.5, compared to the autosomal expectation of close to 0. We then identified putatively sex-linked variants that were heterozygous in the father, homozygous in the mother and where the X-specific allele has a frequency between 0.4 - 0.8 in the male sequencing pool, and >0.98 in the female pool. These thresholds were chosen based on plotting male vs female pool frequencies (see Fig. S2 and JN_05, Appendix 1).

### Identifying the origin of AmhY in C. inconstans

To identify the origin of *Amh* duplication in brook stickleback, we compared the *C. inconstans* copies of this gene to each other and to its orthologs from other stickleback species. It was first necessary to identify *Amh* sequences for available closely related species. To do this we capitalised on already published whole genome sequencing datasets for 7 other stickleback species (*G. aculeatus*: 4 males, 4 females (White *et al*., 2015), *G. nipponicus*: 5 males, 5 females (Yoshida *et al*., 2014), *G. wheatlandi*: 4 males and 4 females (Liu *et al*., 2021), *P. pungitius*: 15 males, 15 females (Dixon *et al*., 2018), *P. tymensis*: 15 males, 11 females (Dixon *et al*., 2018), *P. sinensis*: 13 males, 9 females (Dixon *et al*., 2018) and *Apeltes quadracus*: 4 males, 4 females (Liu *et al*., 2021)).

We aligned adapter and quality trimmed reads from each species to the latest *G. aculeatus* genome assembly, which includes the Y chromosome (Peichel *et al*., 2020). We chose this reference over the *P. pungius* assembly as it is already known that an *Amh* duplicate exists on the assembled Y chromosome of *G. aculeatus* (Peichel *et al*., 2020). Before aligning raw reads, we removed the Y chromosome scaffold from this assembly to ensure that reads from any and all copies of *Amh* in each species would align to the ancestral *Amh* copy on Chromosome 08 in the *G. aculeatus* assembly.

Alignments were again performed using BWA-mem v0.7.17. Reads aligning to *Amh* on *G. aculeatus* Chromosome 08 were subsetted and bcftools v1.10 was used to call variants. Variants were filtered using VCFtools to ensure that each genotype was based on a minimum read depth of 5, had a minimum genotype quality score of 30 and that data for each locus was present in at least 70% of samples within a species dataset. The bcftools consensus tool was then used to produce a consensus sequence for each species using the major (highest frequency) allele at any polymorphic positions. Finally, in species where a sex-linked copy of *Amh* exists, the consensus for the Y copy of *Amh* was output using a custom python script to phase SNPs based on their sex linkage (JN_06, Appendix 1). The resulting consensus sequences of *Amh* and *AmhY* were then aligned using the ClustalW algorithm implemented in MEGA v10.1 (Kumar *et al*., 2018), and a maximum likelihood tree was constructed using a Tamura-Nei nucleotide substitution model with 500 bootstrap replicates, again implemented in MEGA.

Lastly, we predicted the effect of mutations between *Amh* paralogs in *Gasterosteus* and *C. inconstans* using Provean v1.1 (Choi, 2012; Choi *et al*., 2012; Choi & Chan, 2015). Provean compares a query protein to dozens of sequences from other taxa (in this case 66) taken from the NCBI protein database and computes a score which quantifies the conservation of each amino acid. As highly conserved amino acids are expected to be of high functional importance, this “Provean score” can be used to classify amino acid changes between two sequences of interest as putatively neutral (Provean score > -2.5) or putatively deleterious (Provean score ≤ -2.5). We compared *Amh08* and *AmhY* in *C. inconstans* and, for reference, we also compared the ancestral *Amh08* and *AmhY* sequences for all *Gasterosteus* species which were reconstructed using GRASP-suite v2020.05.05.

## Results

### Identification of sex-linked regions in C. inconstans

Comparison of sequencing read depth between males and females failed to identify any large region of the genome with a reduction of read depth in one sex that would be indicative of well-differentiated sex chromosomes. This analysis did, however, identify several extremely narrow regions throughout the genome with either male or female coverage bias (Fig. 1).

**Figure 1.**
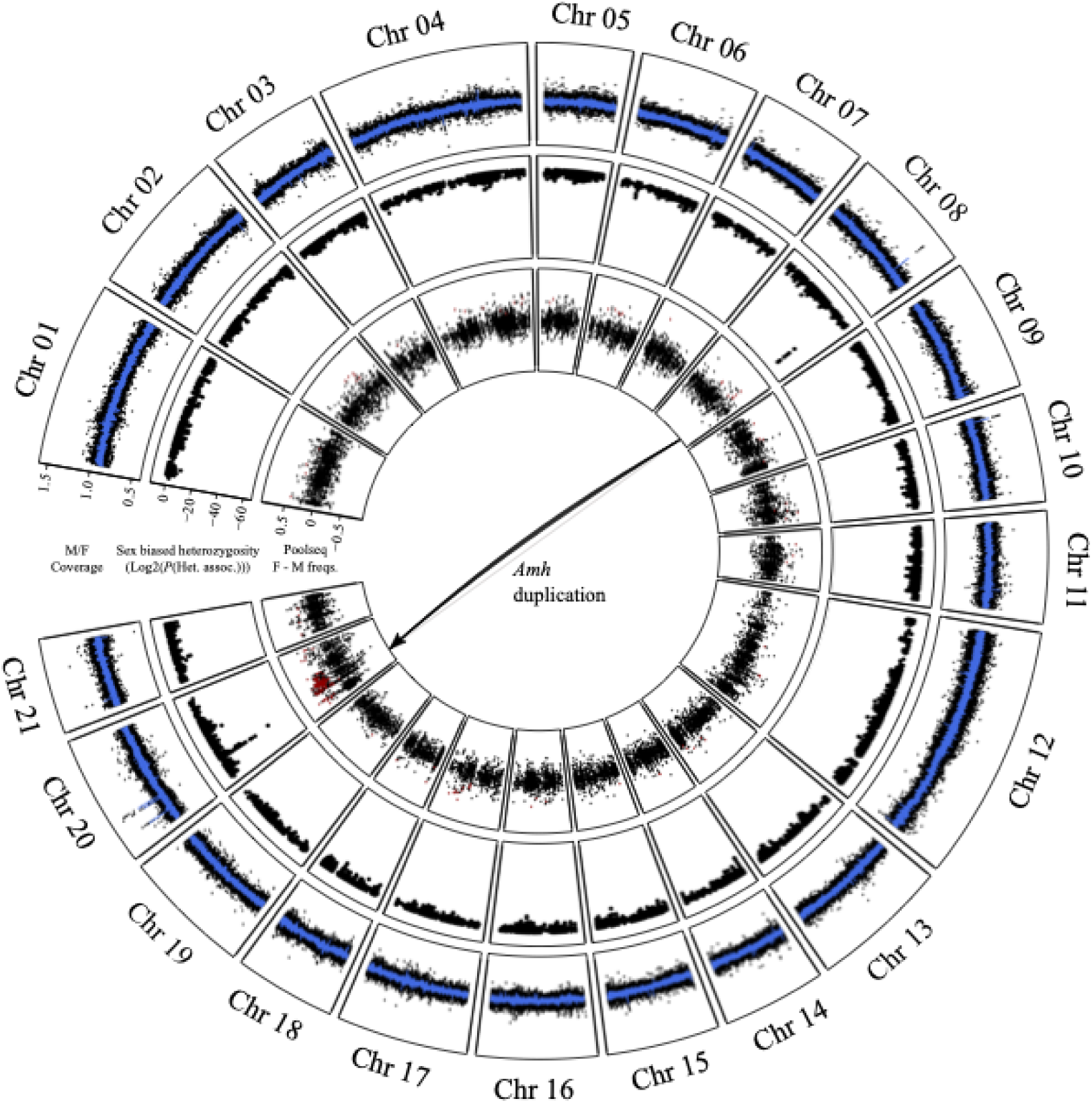
Sex linkage in *C. inconstans*, visualised on the *P. pungitius* genome assembly. Outer track: male / female coverage in 1kb windows. The blue line represents the rolling average across 10 windows. Middle track: results of the test for association of heterozygosity patterns with sex. Inner track: female pool - male pool allele frequencies from the lab cross. The red points represent loci with parental genotypes and pool frequencies that fit expectations of sex linkage (see Methods).

Two such regions were of particular interest. Firstly, on Chromosome 08 there is a clear peak of high male vs female coverage centered at position 16.8-16.9Mb, which exactly matches the position of the gene *Amh* (Fig. S3). Secondly, three peaks of high male vs female coverage co-localise within a ∼5Mb region on Chromosome 20, between position 2-6Mb (Fig. 2). Such peaks of male biased coverage are suggestive of male-specific (i.e. Y chromosome-specific) duplications of these loci.

**Figure 2.**
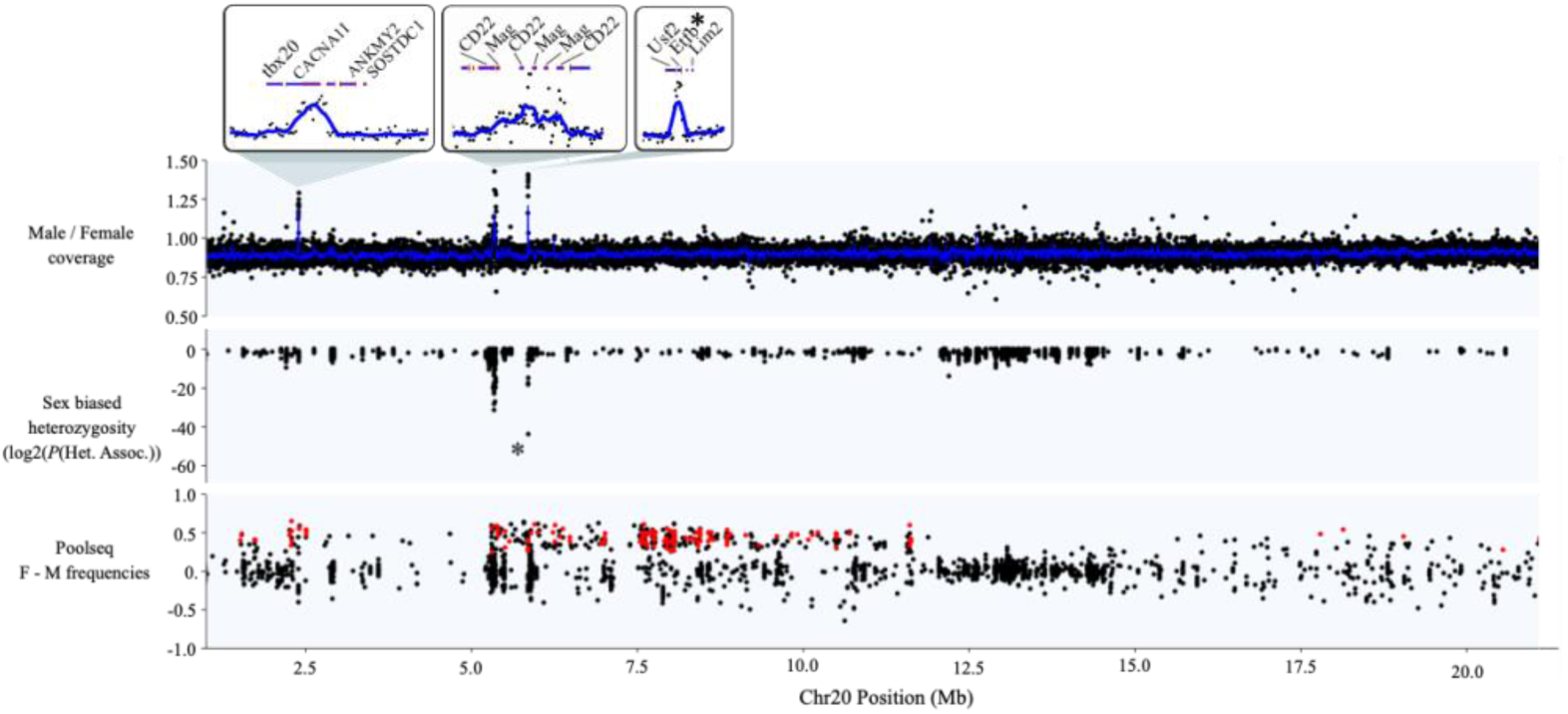
Signals of sex linkage along Chromosome 20, the putative sex chromosome in *C. inconstans*. For the coverage panel, each point represents male / female coverage in a 1kb window. The blue line represents the rolling average across 10 windows. The asterisk in the heterozygosity panel and the zoomed box above represents the single completely sex linked variant that aligned to Chromosome 20 (in the gene *Eftb*). For the pooled sequencing panel, red points represent those with parental genotypes and pool frequencies that fit expectations of sex linkage (see Methods).

The test for the association of heterozygosity patterns with sex was effective at identifying regions of sex linkage and overcoming the excess heterozygosity in the dataset. The vast majority of loci showed high p-values indicative of no sex linkage (Fig. S4). Patterns of sex linkage were localised to two specific regions of the genome (Fig. 1). Of the 10 loci that showed complete sex linkage (i.e. homozygous in all females and heterozygous in all, or all but a few, males), nine of them aligned to a narrow region on Chromosome 08. This region exactly coincides with a peak of male vs. female read depth mentioned above at the location of *Amh* (Fig. S3). The vast majority of the remaining loci showing signs of sex linkage (including one completely sex linked locus) aligned to a ∼0.6Mb region on Chromosome 20 (5.3-5.9Mb), again coinciding with two peaks of increased male vs. female read depth (Fig. 2).

Together, the higher male coverage and the male-biased heterozygosity suggest that a male specific (i.e. Y-specific) *Amh* copy exists in the *C. inconstans* genome (henceforth referred to as *AmhY*) in addition to the ancestral Chromosome 08 copy (henceforth *Amh08*). If this is true, then the coverage of the male specific alleles identified by our heterozygosity analysis, which must have arisen on *AmhY* since the duplication, should be close to ½ that of *Amh08* alleles. Read depth ratios for the nine completely sex linked SNPs showed a clear departure from a 1:1 ratio, and were close to the 1:2 expected ratio, supporting the hypothesis that a Y-specific *Amh* duplicate exists and harbours sex linked variation (Fig. S5). This is the only gene in the genome to show complete sex linkage and is thus a strong candidate for the master sex determination gene in *C. inconstans*.

### Chromosome 20 is the candidate sex chromosome in C. inconstans

While *Amh08* was the only gene in the entire genome to show complete sex linkage, a ∼0.6 Mb region on Chromosome 20 also showed partial sex linkage in our heterozygosity analysis, suggesting that *AmhY* resides somewhere in or near to this region (Fig. 2). This raises the hypothesis that Chromosome 20 is the sex chromosome in *C. inconstans*. Additional support for this hypothesis comes from the fact that one of the 10 loci showing complete sex linkage aligned to this region of Chromosome 20 (specifically within an intron of the gene *Etfb*, which also shows male biased coverage indicative of a Y-specific duplication event). However, our structural variant analysis failed to identify any evidence of the theorised *Amh* duplication to this region, and further, there was not a single structural variant that showed the patterns of sex linkage expected for a Y-specific translocation (Fig. S6). Similarly, our *de novo* assembly of raw sequence reads from 10 males yielded only a single contig containing sequence homologous to *Amh* and this contig also contained regions homologous to the sequence flanking *Amh08*. Contigs aligning to the sex linked region of chromosome 20 were fragmented and showed no sign of containing *AmhY*. We were thus unable to show direct evidence of the theorised translocation event.

However, our analysis of the pooled sequencing data from a laboratory cross yielded 242 putatively sex-linked markers, 158 of which aligned around the previously identified sex-linked region of Chromosome 20 (see red points in Fig. 2). In addition, there are many more markers (plotted in black) in this region with differences in female and male allele frequencies close to 0.5, as expected for sex-linked loci, but which were not heterozygous in the male sample. These are likely also sex linked, but lack heterozygous calls in the father due to allele dropout in low coverage regions. Importantly these results expanded the sex linked region of this chromosome from ∼0.6Mb (from Shunda Lake data alone) to ∼11Mb (between positions 1-12Mb).

### Convergent duplication and recruitment of Amh as the sex determination gene

Consistent with the presence of only four SNPs between *Amh08* and *AmhY* in *C. inconstans*, the phylogenetic analyses of *Amh* sequences from all stickleback confidently clustered *Amh08* and *AmhY* from *C. inconstans* together (Fig. 3). Similarly the *Gasterosteus AmhY* sequences clustered together as an outgroup of the *Gasterosteus Amh08* sequences. These data, therefore, support an independent duplication of *Amh* in *C. inconstans*.

**Figure 3.**
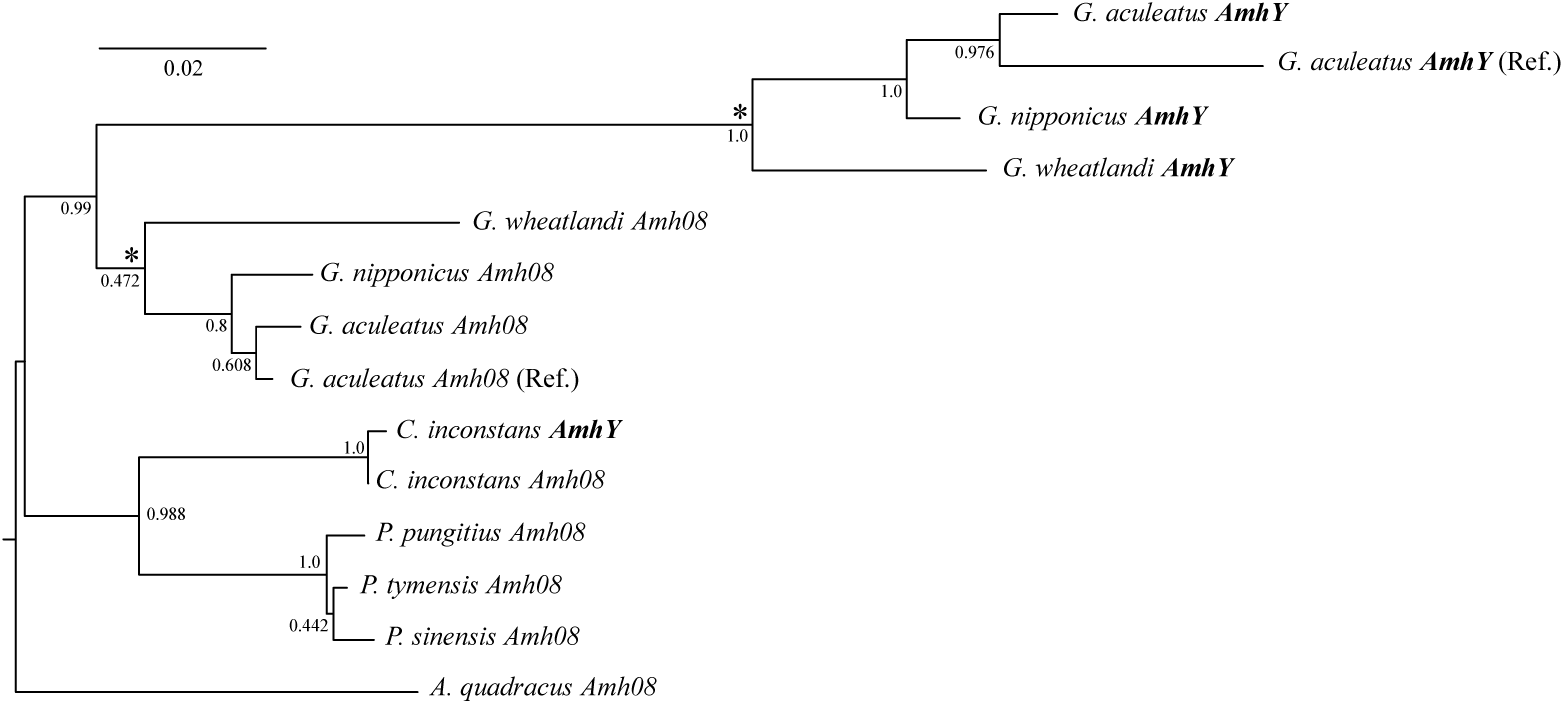
Maximum likelihood phylogeny of *Amh* consensus sequences for eight stickleback species. Node values represent confidence based on 500 bootstraps. Also included are the phased reference *Amh* and *AmhY* sequences from the *G. aculeatus* genome assembly (Peichel *et al*., 2020). Asterisks denote the nodes for which we reconstructed ancestral *Amh* and *AmhY* sequences for our mutation function predictions (see Methods).

Provean analyses of amino acid conservation within *Amh* predicted that of the 93 inferred amino acid changes between the ancestral *Amh08* and *AmhY* sequences in *Gasterosteus*, seven of them are likely to cause deleterious functional changes (Table S1). In contrast, all three of the amino acid changes between *Amh08* and *AmhY* in *C. inconstans* were predicted to be neutral.

## Discussion

The study of taxa with dynamic sex chromosome systems is key to understanding which forces and mechanisms shape the diversity of sex chromosomes and sex determination across the tree of life. In this study we have identified the sex chromosome in *C. inconstans*, a species for which there was previously no information. Thus, there are now five stickleback species with known sex chromosome systems, further solidifying this clade as a valuable model for the study of sex chromosome evolution. Importantly, several attributes of the *C. inconstans* sex chromosome system allow us to speculate on the evolutionary processes at work in this clade, which we discuss in detail below.

### A duplicate of Amh is the candidate master sex determination gene in C. inconstans

Our genomic analyses in *C. inconstans* strongly suggest that the autosomal gene *Amh* has duplicated and that this duplicate has been recruited as the master sex determining gene in this species. However, we can not formally confirm this here. Formal functional validation of this candidate gene (e.g. using transgenics) is beyond the scope of this paper, and our attempts to find direct evidence of the location of *AmhY* on Chromosome 20 failed. However this is perhaps not surprising. There are very few differences between *Amh08* and *AmhY*, thus our assembly approach likely had insufficient unique sequence variation between *Amh08* and *AmhY* to construct different contigs for each. Further, our structural variant analyses suffered from the short insert sizes (300-500 bp) of the sequencing performed here. Such analyses rely heavily on information from the discordant mapping of reads from the same read pairs, and the chances of reads aligning across a structural variant break point greatly decrease with smaller insert sizes.

Nevertheless, the fact that *Amh* was the only gene in the genome to show absolute sex linkage in our population genetics analysis is very strong support for its role as the master sex determination gene. Furthermore, duplications of *Amh* have previously been implicated in sex determination in many fish species, including the pejerreys *Odontesthes hatcheri* and *Odontesthes bonariensis* (Hattori *et al*., 2012; Yamamoto *et al*., 2014), Nile tilapia *Oreochromis niloticus* (Eshel *et al*., 2014; Li *et al*., 2015), lingcod *Ophiodon elongatus* (Rondeau *et al*., 2016), the cobaltcap silverside *Hypoatherina tsurugae* (Bej *et al*., 2017), northern pike *Esox lucius* (Pan *et al*., 2019), Sebastes rockfish *Sebastes schlegelii* (Song *et al*., 2021) and the *Gasterosteus* clade of stickleback (Peichel *et al*., 2020; Sardell *et al*., 2021). *Amh* is also likely used for sex determination in Monotremes, though in this case, both X and Y *Amh* homologs exist and no duplication is apparent. Thus, *Amh* is clearly predisposed to becoming a master sex determination gene in teleosts, as exemplified in the results of the present study by its independent recruitment in two stickleback lineages within the last 25-30 My.

It is interesting to note that, with the exception of monotremes, it is always a duplicate of *Amh* that determines sex, with no examples, to our knowledge, of the autosomal copy being recruited for sex determination in teleosts. This suggests that duplication is an important process in the recruitment of this gene as the master sex determination gene and begs the question as to why that may be. One hypothesis is that the ancestral, autosomal copies of *Amh* in these species play some vital role that cannot be altered but can be circumvented via its duplication and subsequent sub- or neo-functionalization. A second hypothesis is that the duplication itself is the sex determining mutation, i.e. simply increasing the dose of this gene is enough to initiate male development. We propose that the second of these hypotheses is more likely, based on two lines of reasoning. The first relies on the observation that, in all of the cases above (which are all XX/XY systems), the duplication is Y-specific and is absent from the X. If the duplicate does not determine sex when it first arises, then it is free to segregate like any autosomal gene on both homologs of its resident chromosome pair and, at least in some cases, it might be fixed. If one allele of the duplicate later acquires the male determination role, an X copy would still exist. Thus, to match the observation that no X homolog exists in any of the eight species discussed here, we would need to invoke multiple losses of the X homologs, which is unlikely. Alternatively, if, from the moment it arose, the duplicate could determine sex, homozygosity would not be possible, as that would require males mating with males. The lack of an X homolog is, therefore, an intrinsic prediction of a scenario where the duplicate *Amh* determines sex from the moment of duplication.

The second line of reasoning rests on our results suggesting that the three amino acid changes between *Amh08* and *AmhY* in *C. inconstans* do not substantially alter the function of the protein. This would imply that no functional change was necessary for *AmhY* to assume the role of sex determination in *C. inconstans* and implicates the increased gene dose as the likely sex determination mechanism. In contrast to the *Amh* duplication in *C. inconstans*, the duplication events in northern pike, *Gasterosteus* stickleback, rockfish, pejerreys, lingcod, the cobaltcap silverside, and Nile tilapia, are relatively old. In all of these cases, there is substantial protein sequence divergence between the ancestral and duplicated *Amh* copies, however it is not possible to infer whether these mutations were important in the recruitment of the *Amh* duplicates for sex determination in these species, or whether they have arisen since. It would be interesting to follow up on this question in future studies, for example, by creating transgenic XX individuals with an additional *Amh* copy to test for the effect of increased *Amh* dose on sex determination, in the absence of any amino acid changes.

Another interesting observation is that all eight of the species known to have independently recruited *Amh* duplicates for sex determination (including *C. inconstans* as shown here) belong to the clade Teleostei. This is unlikely to be purely coincidental. In most vertebrates, *Amh* (Anti-Müllerian hormone) is responsible for inhibiting the development of the female reproductive tract (Müllerian ducts) during embryogenesis (Capel, 2017) and thus promoting male development. However, teleosts lack Müllerian ducts (Adolfi *et al*., 2019). The exact role of *Amh* in teleosts is not yet well understood, however gene expression data from Nile tilapia (Li *et al*., 2015) and pejerreys (Hattori *et al*., 2012; Yamamoto *et al*., 2014) shows expression of the *Amh* duplicate genes occurs just prior to gonadal differentiation, suggesting that they likely play an important role in the proliferation and differentiation of germ cells during gonad development. Based on the above, one could speculate that the loss of Müllerian ducts in teleosts has freed *Amh* of its primary role of Müllerian duct suppression and this allows it to be more easily repurposed for sex determination.

More broadly, while several theoretical studies have considered the evolutionary forces that might drive a new MSD to fixation, one understudied component of sex chromosome turnovers is the rate at which alternative MSDs arise. Given that the genetic architecture of sex determination pathways differ drastically among taxa (Capel, 2017), it is not unreasonable to expect that taxa also differ in the number of possible alternative MSDs that exist. Thus, it is possible that in some lineages, the rate of sex chromosome turnovers is limited by the constraints of their existing sex determination pathway and the frequency with which alternative MSDs can arise, while, in others, there may be numerous potential MSDs, which can readily evolve via simple mutations (e.g. gene duplication). The *Amh*/*AmhrII* pathway in teleosts may be an example of such a scenario, and may in turn help explain their rapid turnover rate. Indeed, *AmhrII*, the receptor of *Amh* has also been found to be the sex determiner in pufferfish (Kamiya *et al*., 2012; Ieda *et al*., 2018), the ayu *Plecoglossus altivelis* (Nakamoto *et al*., 2021) and the yellow perch *Perca flavescens* (Feron *et al*., 2020).

### Sex chromosome turnover in Stickleback

With the results of this study, there are now three independently evolved sex chromosome systems known in sticklebacks, Chromosome 19 (*AmhY*) in *Gasterosteus* (with a further two independent Y-autosome fusion events within this clade), Chromosome 12 in *Pungitius pungitius* and now Chromosome 20 in *C. inconstans*. There must therefore be a minimum of two sex chromosome turnover events among these species. However, the inability to find sex linkage signal on any of these chromosomes in *P. tymensis* or *P. sinensis* (Dixon *et al*., 2018) or in *A. quadracu*s (Ross et al. 2009) might be suggestive of more turnovers.

The sex chromosome of *C. inconstans* shows the lowest divergence of any of the sex chromosomes now described in stickleback when compared to the heteromorphism observed in *Gasterosteus* (Ross & Peichel, 2008; Peichel *et al*., 2020; Sardell *et al*., 2021) and the large region of differentiation found in *P. pungitius* (Dixon *et al*., 2018). Consistent with a lack of extensive degeneration on the Y, there is no evidence of a reduction in read coverage in males relative to females. Furthermore, there are very few loci that show evidence of differentiation between the X and the Y (i.e. differences in heterozygosity between males and females). In fact, if we are correct in our hypothesis that *AmhY* lies near to the gene *Eftb* on Chromosome 20, then the completely sex-linked region of this chromosome may be on the order of 1Mb in length. Given that, in general, recombination loss and sex chromosome differentiation expands outwards from the sex determination locus over time, this would suggest that the turnover event in *C. inconstans* was very recent, and that the sex chromosomes in this species are young. Unfortunately, we cannot precisely estimate the size and level of differentiation of the sex-linked region without a good quality long read assembly of the X and Y chromosomes of *C. inconstans*. However, if it is recent, as our current data suggests, it may be possible to infer the identity of the ancestral sex chromosome pair by searching for signals left behind during its time in this role (e.g. reduced effective population size, increased repeat content (Vicoso & Bachtrog, 2013)). This would be an interesting topic of further study and could help to further characterise the transitions among sex chromosomes in stickleback.

Interestingly, our coverage analyses also identified several other genes in this sex-linked region which seem to have have male specific copies (see inlays in Fig. 2). None of these genes have roles that have previously been associated with sex, though it is possible that their duplication and linkage with the sex determination gene may still be adaptive, for example, as a means of resolving genomic conflict at a sexually antagonistic locus (Bergero *et al*., 2019). The gene locations in this study are based on those in the *P. pungitius* assembly, however, given that they show evidence of sex linkage, its is likely that these genes are in approximately the same location in *C. inconstans*. In addition, a manual examination of the *G. aculeatus* genome assembly places *tbx20* and *ANKMY2* next to *Mag* between positions 5 - 6 Mb on Chromosome 20 (*CACNA1I, CD22, Eftb*, and *Lim2* were unfortunately not annotated in *G. aculeatus*). These locations are, therefore, likely ancestral. However, given the proximity of *tbx20, ANKMY2*, and *Mag* in *G. aculeatus*, there may, in fact, be an inversion around 2 - 5 Mb specific to *P. pungitius*, which would explain the distance between the sex linked *tbx20* duplicate and the region of sex linkage identified in the wild-caught data and the dearth of sex-linked variants in the lab cross in this region. Thus, it seems this region may be particularly prone to structural variation. Again, a high quality reference assembly for the *C. inconstans* X and Y chromosomes is needed to resolve the speculation above.

In the context of studying young sex chromosomes, it is useful to highlight the utility of different data types in our analyses. Though pooled sequencing strategies lose individual haplotype information and the ability to examine heterozygosities, the fact that this data came from an F1 cross limited the number of recombination events in the dataset. The resulting large blocks of linkage disequilibrium along the genome made this dataset ideal for looking for a broad signal of sex linkage and, in this case, was essential for confidently identifying the sex chromosome. In fact, given how broad the sex linkage signal is in this cross, individual based sequencing would have been overkill, as increasing marker density far beyond the size of linkage blocks would add no more biological information. It should be noted, however, that such an approach will always overestimate the extent of true recombination suppression between sex chromosomes because the detection of rare recombination events is limited by the number of individuals in the cross. In contrast, the population level dataset from Shunda Lake represents a large sampling and sequencing effort in terms of both money and time, but provided the extremely fine resolution needed to distinguish between the complete sex linkage of a single gene, infer its duplication, and to infer partial sex linkage elsewhere in the genome. Importantly, this resolution comes not only from the high marker density and numerous samples, but from the large number of recombination events that have happened in the ancestors of all individuals sampled. Incorporating long coalescent times into a sample set captures linkage information from thousands of recombination events, and it is this that allowed us to so finely map sex-linked regions of the genome in this study. We highlight this point with the hope of aiding future researchers to design the most informative and cost effective dataset for their purposes.

## Data availability

The sequencing data in this study is in the process of being uploaded to the SRA, accessions will be included here before publication. All custom python scripts can be found in Jupyter notebooks in the appendix, along with relevant intermediate files from the evolutionary analyses.

## Supplementary figures

**Figure S1.**
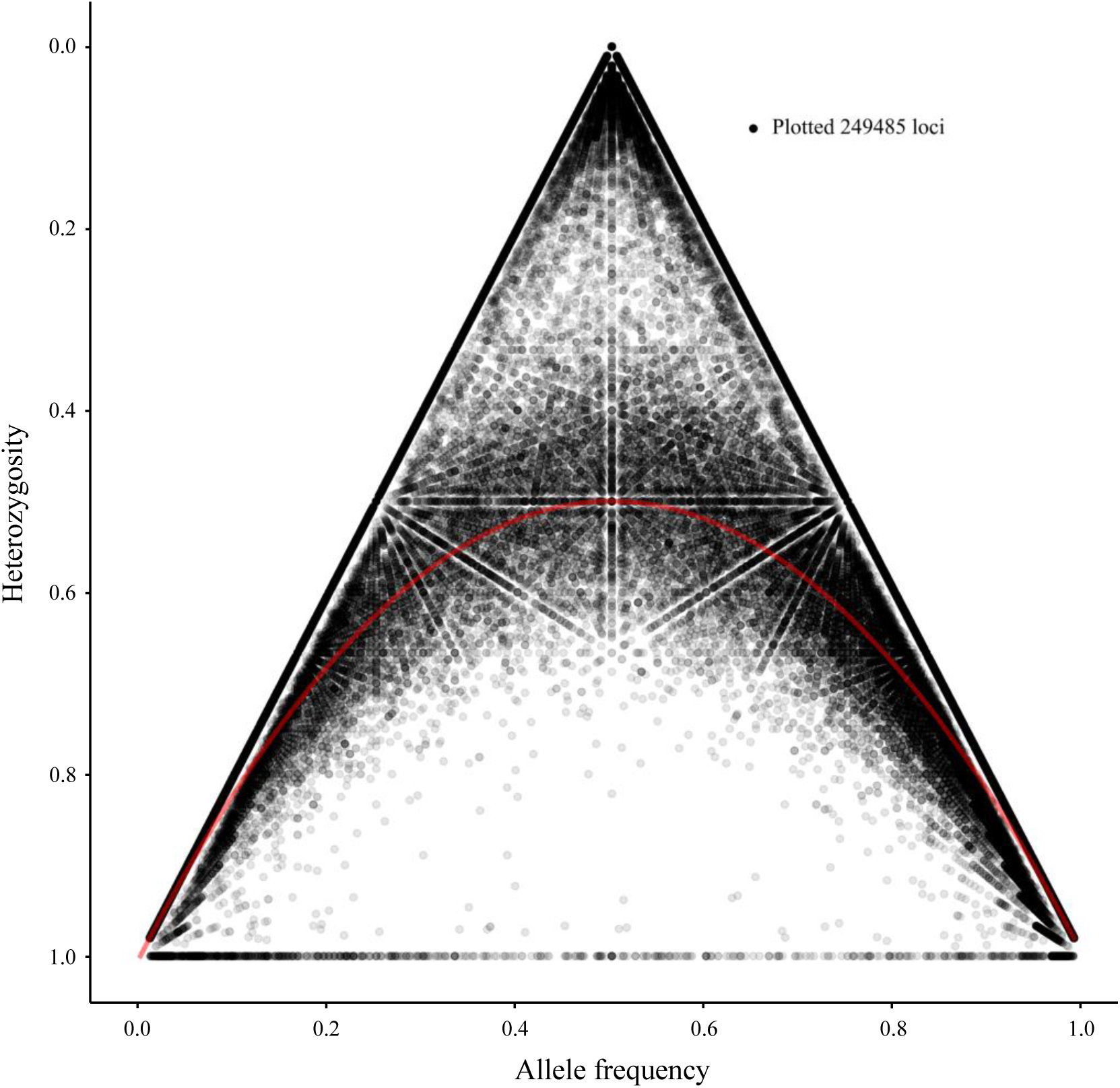
Comparison of allele frequency and heterozygosity to assess the quality of post-filtering genotype calls in the Shunda Lake whole genome resequencing dataset. The red line represents the expectation under Hardy-Weinberg equilibrium.

**Figure S2.**
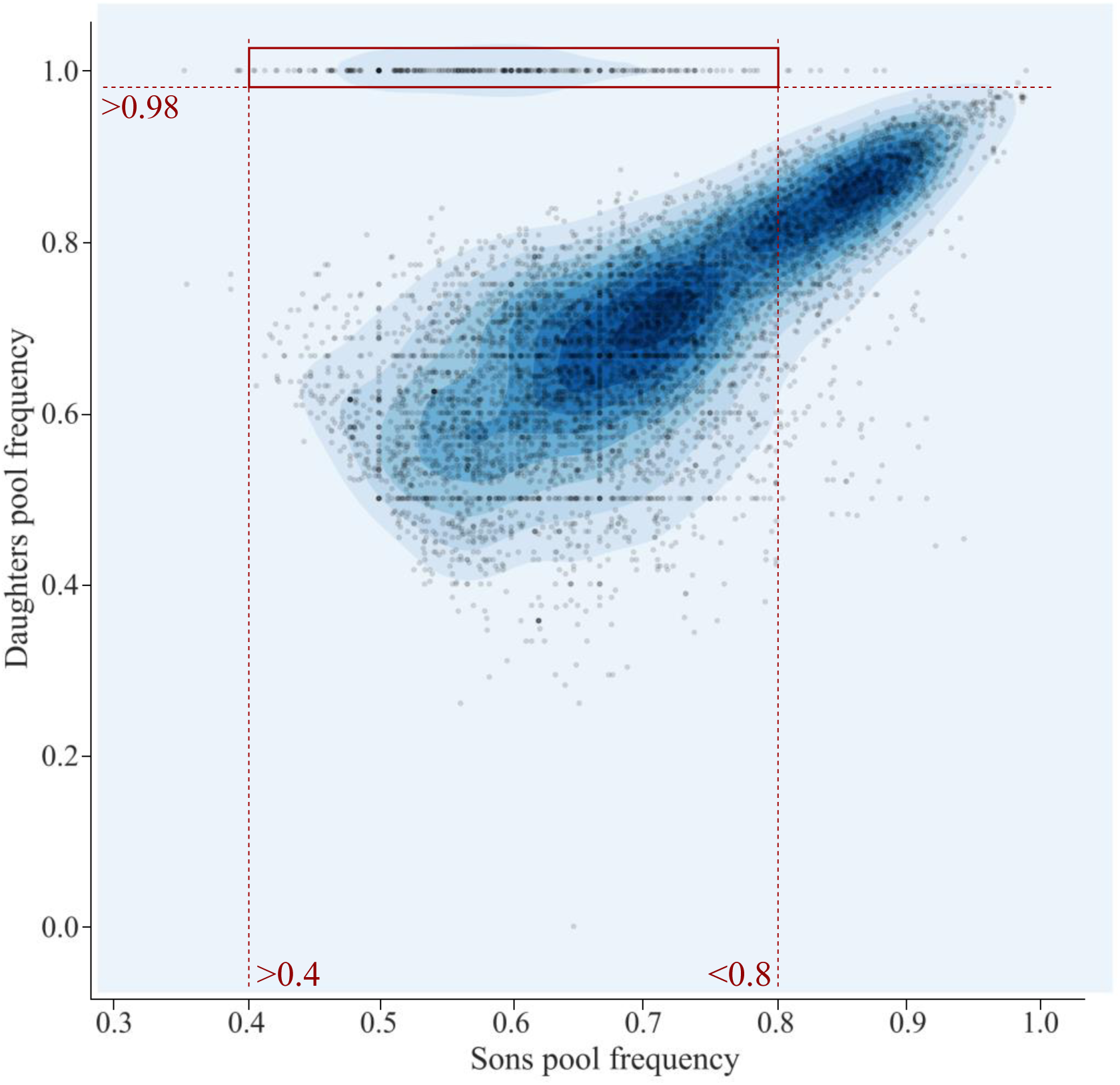
Scatter and density plot comparing the frequency of alleles in the pooled sequencing of sons and daughters for loci that are heterozygous in the father and homozygous in the mother. The red box shows the cut-offs used to label points as putatively sex linked in figures 1 and 2.

**Figure S3.**
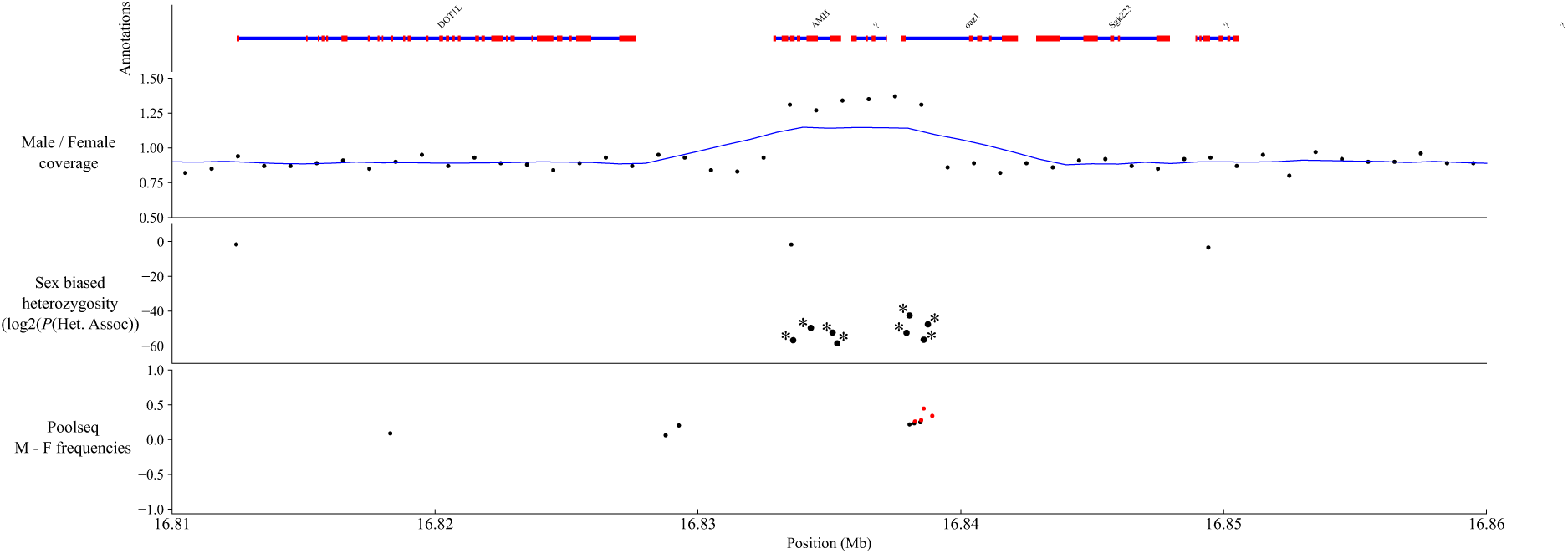
Zoomed in view of sex linkage signal around *Amh08* in *C. inconstans*. An increase in male to female coverage suggests a male specific copy of *Amh* exists. Blue line represents the rolling average across 10 1kb windows. Asterisks on the heterozygosity association panel represent eight completely sex-linked variants.

**Figure S4.**
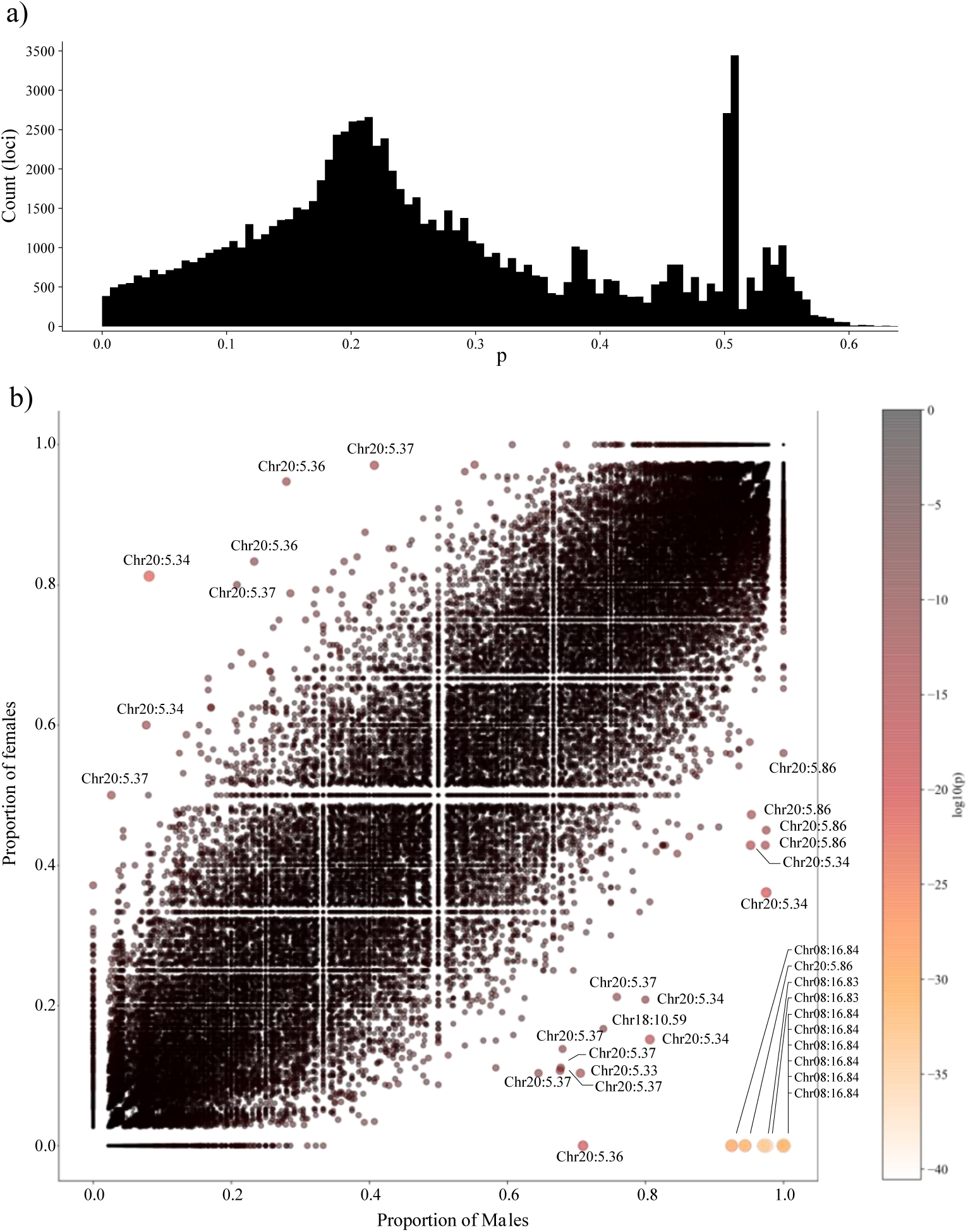
a) Distribution of *P* values for association between sex and heterozygosity in the Shunda lake whole genome resequencing dataset. b) Scatter plot showing proportion of heterozygous males vs proportion of heterozygous females for each SNP. Points are coloured and sized according to the *P* value for the association between heterozygosity and sex. A subset of the most sex-linked loci are labelled to illustrate the enrichment of loci on Chromosomes 08 and 20.

**Figure S5.**
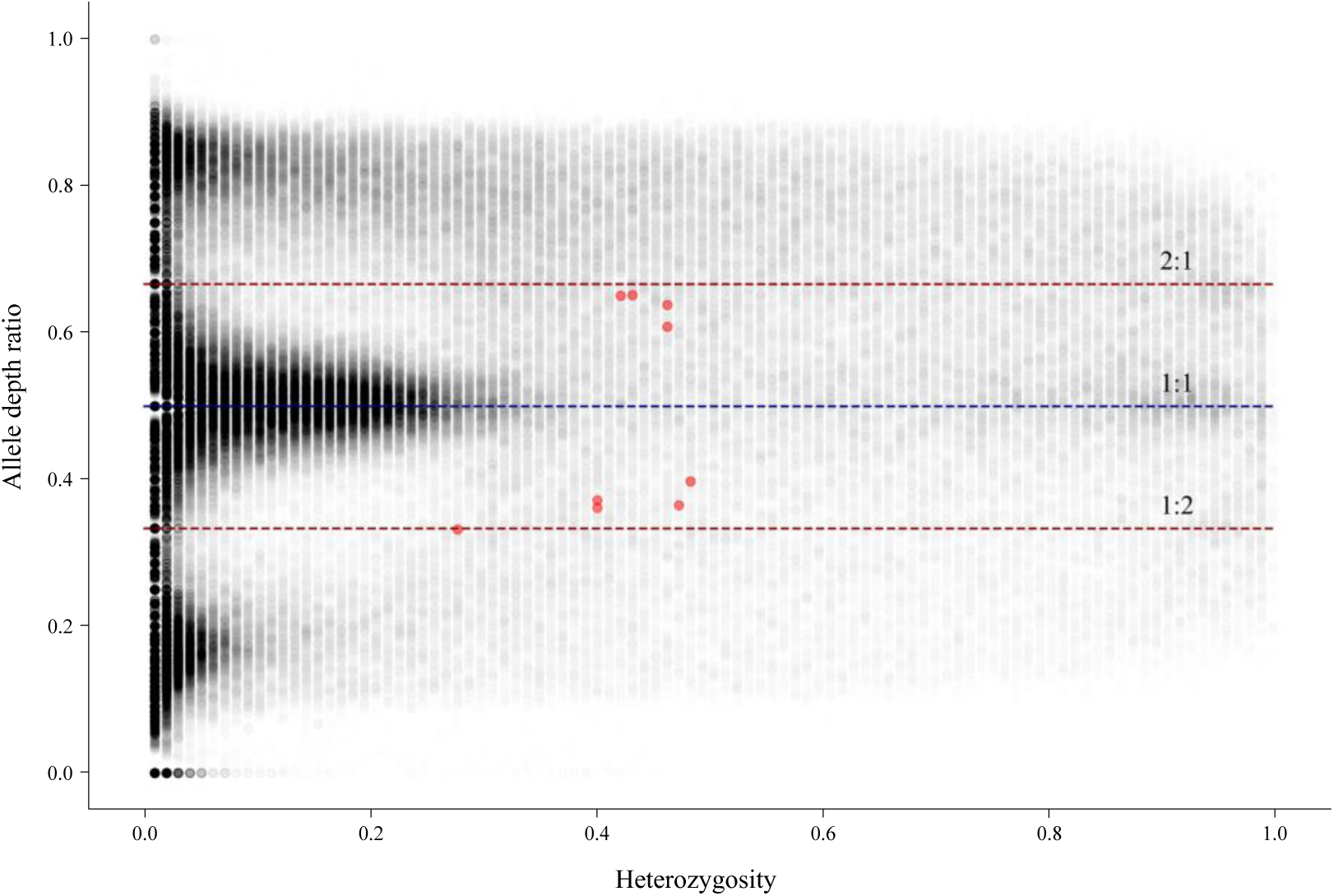
Comparison of heterozygosity and allelic depth ratio for each variant in the Shunda Lake whole genome resequencing dataset, as calculated by HDplot. Red dashed lines represent the allele depth ratio expected for loci with a duplicated sex linked copy. Red points represent the 10 completely sex-linked markers in the dataset.

**Figure S6.**
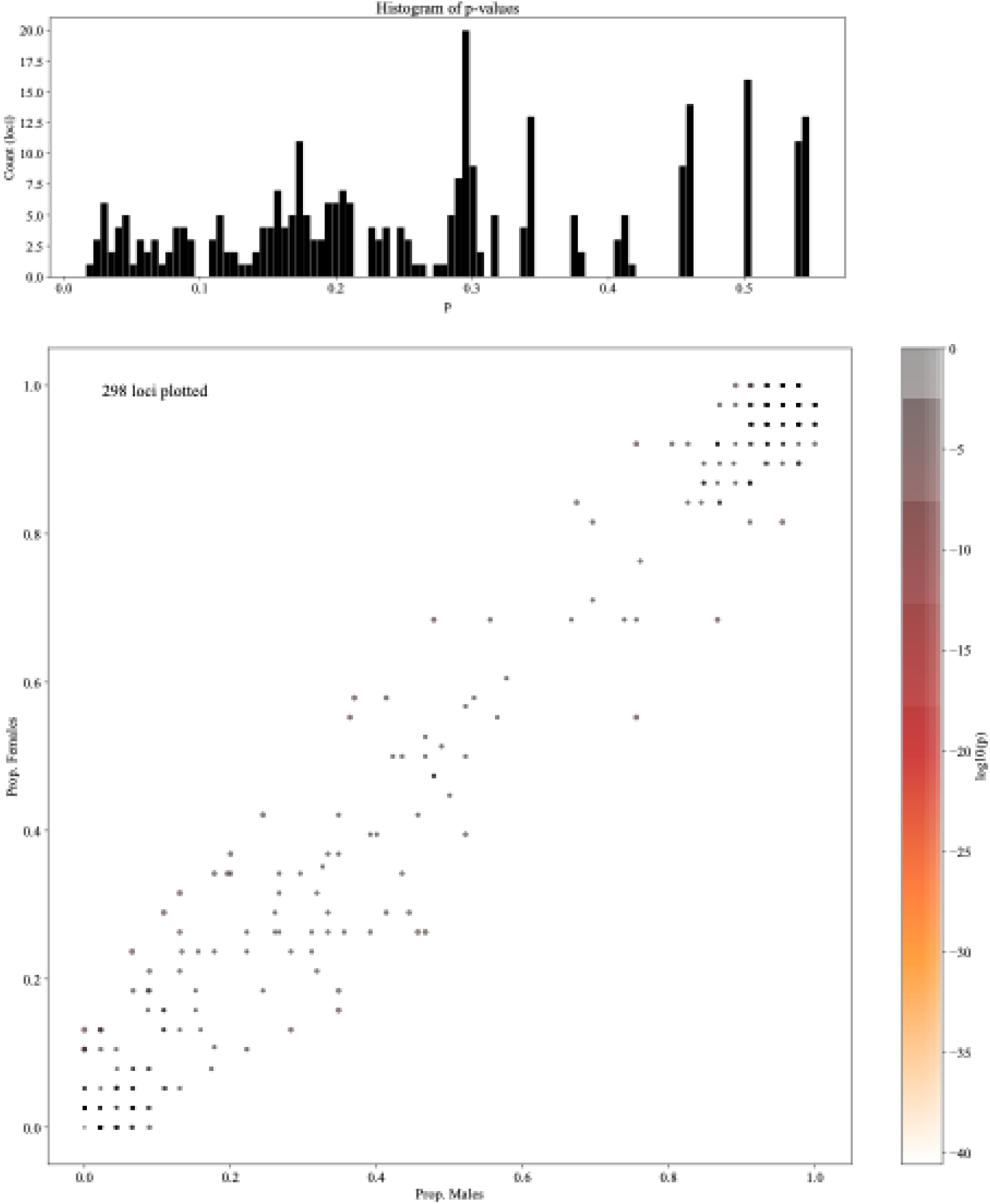
Test for association between sex and heterozygosity of structural variants called using DELLY. Points in the scatter plot are coloured and sized based on their *P* value (scale is identical to Fig. S4b). No variant shows any sign of association of sex linkage.

**Supplementary Table S1.**
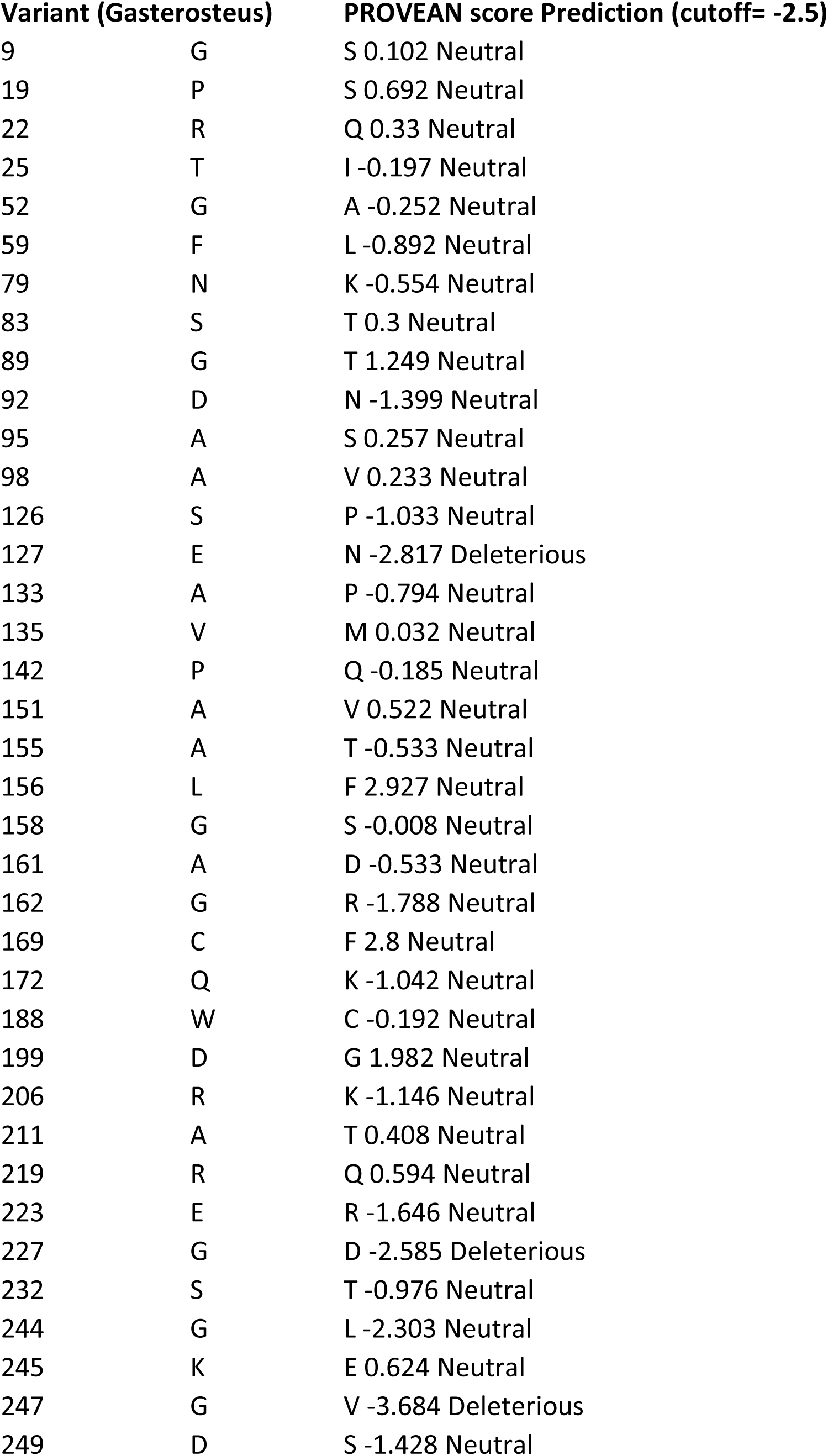
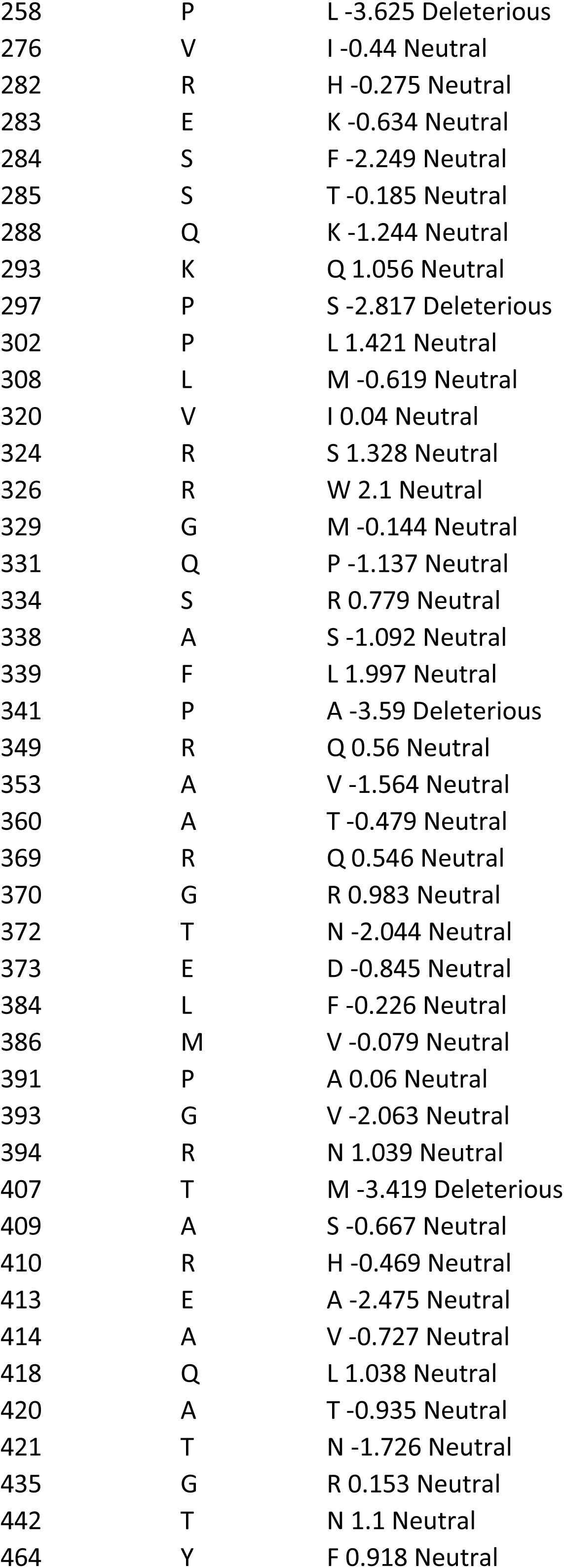
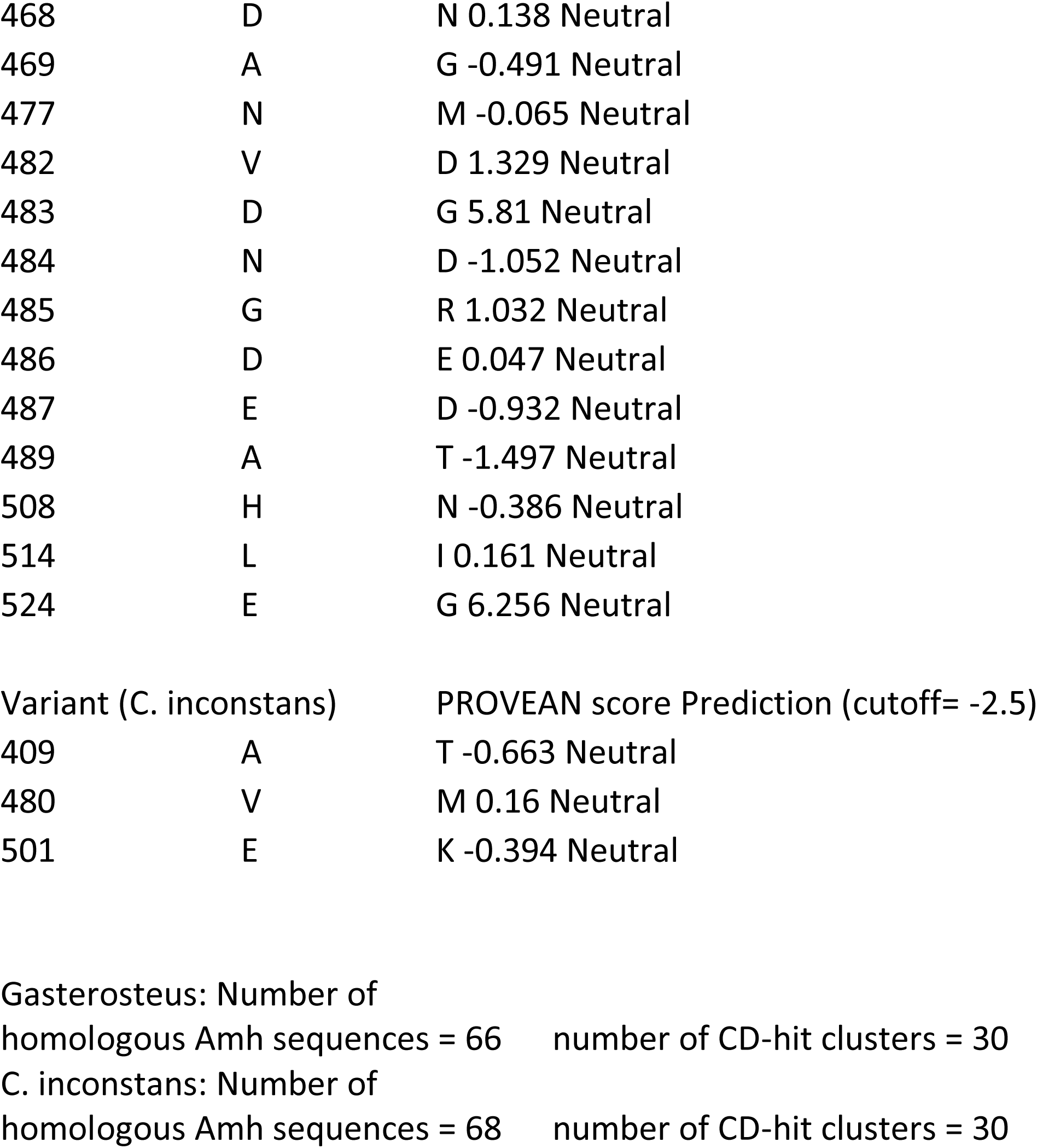
Outputs from the Provean predictions for the impact of amino acid changes among *Amh* paralogs. Variants are listed in the format: Position, *Amh08* amino acid, *AmhY* amino acid.

## Notes

### Competing Interest Statement

The authors have declared no competing interest.

